# Selective depletion of AMH-expressing granulosa cells *in vivo* impairs follicular development and fertility in female mice

**DOI:** 10.64898/2026.08.04.742350

**Authors:** Tsutomu Endo, Mayu Tamemasa, Kana Hayakawa, Fuka Okada, Natsumi Oyama, Kyoko Watanabe, Tianfu Lai, Yuki Nakano, Yoshifumi Fujioka, Motohito Goto, Riichi Takahashi, Ayako Tomita, Koji Sugiura, Yoshikazu Hirate, Naoaki Mizuno, Yoshiakira Kanai, Masami Kanai-Azuma

**Affiliations:** Department of Experimental Animal Model for Human Disease, Graduate School of Medical and Dental Sciences, Institute of Science Tokyo, Tokyo, Japan; Center for Experimental Animals, Institute of Science Tokyo, Tokyo, Japan; Department of Animal Resource Sciences, Graduate School of Agricultural and Life Sciences, The University of Tokyo, Tokyo, Japan; Department of Veterinary Anatomy, Graduate School of Agricultural and Life Sciences, The University of Tokyo, Tokyo, Japan; Central Institute for Experimental Medicine and Life Science (CIEM), Kanagawa, Japan

**Keywords:** AMH, granulosa cells, folliculogenesis, fertility

## Abstract

In mammals, ovarian follicle development is a highly coordinated process that underlies female fertility. Granulosa cells expressing anti-Müllerian hormone (AMH) are widely used as a marker of growing follicles. However, the *in vivo* roles of granulosa cells in follicular development and female fertility remain unclear. Here, we analyzed AMH-toxin receptor-mediated cell knockout (AMH-TRECK) transgenic (Tg) mice on a NOG background, in which AMH-expressing granulosa cells are specifically depleted by diphtheria toxin (DT). We first found that, after a single DT injection into postnatal AMH-TRECK Tg females, AMH-expressing granulosa cells in primary and secondary follicles exhibited cleaved caspase-3 signals 1 day later and were depleted 4 days later. Second, after repeated DT injections weekly from 1 to 7 weeks of age in AMH-TRECK Tg females, antral follicles and corpora lutea were rarely observed, and the numbers of primordial, primary, and secondary follicles were decreased. Following PMSG and hCG stimulation, repeated DT-injected Tg females exhibited a reduced number of ovulated oocytes with a low proportion of mature oocytes, resulting in reduced IVF rates and fertility. Further, after a cessation of repeated DT treatment, ovarian weight and follicular development recovered: the numbers of primary, secondary, and antral follicles were recovered, whereas the primordial follicle pool remains reduced. We conclude that selective depletion of AMH-expressing granulosa cells *in vivo* impairs follicular development and fertility. Our model enables assessment of the *in vivo* effects of granulosa cell depletion and may provide a useful platform for future transplantation-based studies to understand complex follicular dynamics.

## Introduction

Ovarian follicular development is a highly coordinated process that underlies female fertility and requires dynamic interactions between oocytes and surrounding somatic cells (granulosa cells). Granulosa cells play essential roles in follicle growth, oocyte quality, and ovulation through the production of hormones and paracrine factors (Matzuk et al., 2002; Richards and Pangas 2010). Investigating the *in vivo* roles of granulosa cells is important for elucidating their contributions to follicular development and female fertility.

During folliclogenesis, ovarian follicles progress through the primordial, primary, secondary, and antral stages before reaching the preovulatory stages (Richards and Pangas 2010). Follicle-stimulating hormone (FSH) promotes the growth of late secondary (as known as preantral) and antral follicles to the preovulatory stage, which is characterized by a large antral cavity and expanded granulosa cell population (McGee and Hsueh 2000; Liu et al., 2002). Following the LH surge, the follicle ruptures to release the oocyte during ovulation, and the remaining follicular cells luteinize to form a corpus luteum. After birth, anti-Müllerian hormone (AMH) is specifically expressed in supporting cells of both testes and ovaries, namely Sertoli cells and granulosa cells, respectively (Lecureuil et al., 2002). In the mouse ovary, AMH is not expressed in granulosa cells of primordial follicles but is first expressed in those of primary follicles on postnatal day 3 (P3) or P4 (Durlinger et al., 2002). AMH expression is high in primary and secondary follicles, decreases in early antral follicles, and is absent in late antral and preovulatory follicles (Visser et al., 2006). Therefore, AMH-expressing granulosa cells are widely used as a marker of growing follicles (Durlinger et al., 2002; Visser et al., 2006).

We previously generated the AMH-toxin receptor-mediated cell knockout (AMH-TRECK) transgenic (Tg) mouse model, in which the human diphtheria toxin (DT) receptor is specifically expressed in AMH-expressing Sertoli and granulosa cells of testes and ovaries, respectively (Shinomura et al., 2014); in this TRECK system (Saito et al., 2001; Furukawa et al., 2006), a single DT injection into postnatal AMH-TRECK Tg mice specifically depletes AMH-expressing Sertoli and granulosa cells. This mouse model enables investigation of the *in vivo* roles of AMH-expressing granulosa cells in follicular development and female fertility. However, the comprehensive effects of AMH-expressing granulosa cell depletion on postnatal and adult ovaries, as well as female fertility, remain unclear.

In the present study, we address whether selective *in vivo* depletion of AMH-expressing granulosa cells affects ovarian development and female fertility using AMH-TRECK Tg mice. Specifically, we first examined the early effects of a single DT injection on postnatal ovaries in AMH-TRECK Tg females. We then performed repeated DT injections into AMH-TRECK Tg females from the postnatal period to adulthood and comprehensively evaluated ovarian morphology, ovulation, *in vitro* fertilization (IVF), preimplantation embryonic development, and *in vivo* fertility. In addition, we examined whether the effects of repeated DT injections on AMH-TRECK Tg ovaries were reversible after cessation of DT injections. All experiments in the present study were conducted using AMH-TRECK Tg mice on a NOG genetic background, generated by backcrossing the previously reported C57BL/6 AMH-TRECK Tg line (Shinomura et al., 2014) onto the NOG background (Ito et al., 2002), given its potential utility for future cell transplantation studies.

## Materials and Methods

### Animals

All animal experiments were approved by the Animal Care and Use Committee of the Central Institute for Experimental Medicine and Life Science (CIEM) and the Institute of Science Tokyo. Mice were maintained in designated animal rooms under a 12-h light–dark cycle with ad libitum access to food and water at CIEM and the Institute of Science Tokyo. NOG mice, originally generated as previously described (Ito et al., 2002), were provided from CIEM. C57BL/6 AMH-TRECK Tg mice (line #94), carrying an *AMH* promoter-driven human diphtheria toxin receptor (DTR; HBEGF-mut)-EGFP transgene (Figure 1A) and previously established as described (Shinomura et al., 2014), were backcrossed with NOG mice for eight generations to generate AMH-TRECK Tg NOG mice.

**Figure 1.**
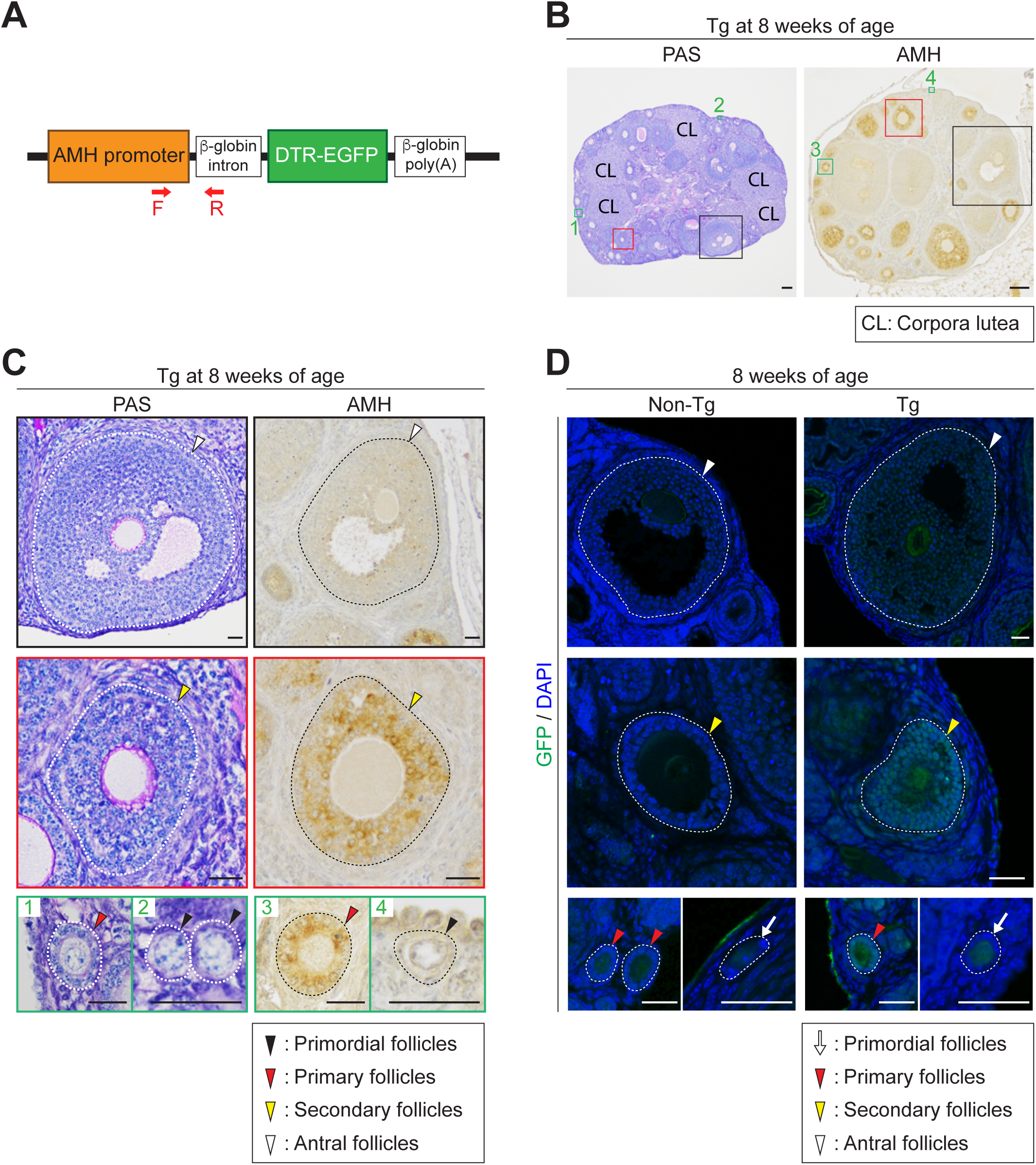
AMH expression in granulosa cells of adult ovarian follicles in AMH-TRECK Tg NOG mice. (A) Schematic representation of the AMH-TRECK transgene. The diphtheria toxin receptor (DTR; HBEGF-mut) is fused to EGFP downstream of the exogenous *AMH* promoter. Red arrows, the forward (F) and reverse (R) primers for genotyping PCR shown in Supplementary Figure S1. (B, C) Ovary sections from Tg females at 8 weeks of age, stained with hematoxylin and periodic acid-Schiff (PAS) (left) or immunostained for AMH (right). CL, corpora lutea. Boxed regions (black, red, and green #1-4) in (A) are enlarged in (B). Arrowheads, primordial (black), primary (red), secondary (yellow), and antral (white) follicles. Dashed lines, basement membranes of follicles. Scale bars, 100 μm in (A) and 25 μm in (B). (D) Ovary sections from non-Tg (left) and Tg (right) females at 8 weeks of age, immunostained for GFP (green) with DAPI counterstain (blue). White arrows, primordia follicles. Arrowheads, primary (red), secondary (yellow), and antral follicles. Dashed lines, basement membranes of follicles. Scale bars, 25 μm.

### Genotyping

Genotyping to identify AMH-TRECK Tg and non-Tg mice was performed by multiplex PCR using primers specific for the AMH*-*TRECK transgene (282 bp) and *Gdnf* (187 bp) as an internal control (Supplementary Figure S1). Primer sequences were as follows: AMH*-*TRECK (forward), 5’-AGAAAGGGCTCTTTGAGAAGGCCACTCTGC-3’; AMH*-*TRECK (reverse), 5’-CCATCCTAAACAACACCCTGAAAACTTTGC-3’; *Gdnf* (forward), 5’-TTAGGTCAGAAAGCGAATCAGAAAAGCCAC-3’; *Gdnf* (reverse), 5’-GGCAGGGGCGAGAAGACAAGCAGCCTGCAC -3’.

### Diphtheria toxin injection

For DT injection experiments, mice at P7 and P14 received subcutaneous injections of DT (322326, Sigma-Aldrich, St Louis, MO, USA) (4 μg/mL in PBS) at 10 μL/g body weight (40 μg/kg body weight), and mice at P21 and older received intraperitoneal injections of DT (4 μg/mL in PBS) at 10 μL/g body weight (40 μg/kg body weight).

### Histology

Ovaries were fixed in Bouin’s solution for 2 hr at room temperature, embedded in paraffin, sectioned at 5 µm thickness, and stained with hematoxylin and periodic acid-Schiff (PAS). All sections were observed under a light microscope.

### Immunostaining

Ovaries were fixed in Bouin’s solution for 2 hr at room temperature or in 4% (w/v) paraformaldehyde (PFA) overnight at 4°C, embedded in paraffin, and sectioned at 5 µm thickness. Slides were de-waxed, rehydrated, and heated in 10 mM sodium citrate buffer (pH 6.0).

For fluorescence detection, slides were then blocked with 2.5% donkey serum in PBS for 30 min at room temperature, incubated with a primary anti-GFP antibody (1:500 dilution; chicken polyclonal, ab-13970, Abcam, Cambridge, UK) overnight at 4°C, washed with PBS, incubated with the secondary donkey anti-chicken IgY (IgG) (H+L) antibody conjugated to Alexa Fluor 488 (1:500 dilution; 703-545-155, Jackson ImmunoResearch, West Grove, PA, USA) for 30 min at room temperature, and washed with PBS. The slides were counterstained and mounted with ibidi Mounting Medium with DAPI (50011, ibidi GmbH, Gräfelfing, Germany), and fluorescence images were acquired using an LSM 700 laser-scanning confocal microscope (Zeiss, Oberkochen, Germany).

For colorimetric detection, slides were then blocked with 2.5% horse serum (Vector Laboratories, Newark, CA, USA) for 30 min, incubated with the primary antibodies for 1 hr, washed with PBS, incubated with the secondary antibodies (ImmPRESS-HRP detection kit, Vector Laboratories) for 30 min, and washed with PBS, at room temperature. The slides were developed using a DAB substrate kit (Vector Laboratories), counterstained with Mayer’s hematoxylin, dehydrated, and mounted with Permount (Fisher Scientific, Waltham, MA, USA). The primary antibodies for colorimetric detection were as follows: anti-AMH (MIS) (1:500 dilution; goat polyclonal, sc-6886, Santa Cruz Biotechnology, Dallas, TX, USA) and anti-cleaved caspase-3 (1:100 dilution; rabbit monoclonal, 9579, Cell Signaling Technology, Danvers, MA, USA).

### Follicle count

Follicle counts were performed on every third 5-µm section containing a visible germinal vesicle (Sugiura et al., 2010). Follicles were classified according to the following criteria: primordial follicles, with a single layer of flattened pregranulosa cells surrounding the oocyte; primary follicles, with a single layer of cuboidal granulosa cells; secondary follicles, with two or more layers of granulosa cells and a theca layer; and antral follicles, with a fluid-filled antrum.

### *In vitro* fertilization and embryo culture

Mouse IVF was performed as described previously (Endo et al., 2024), with minor modifications. In brief, 8-week-old females were given an injection of 7.5 IU of pregnant mare serum gonadotropin (PMSG; ASKA Pharmaceutical Co., Tokyo, Japan), followed 48 hr later by 7.5 IU of human chorionic gonadotropin (hCG; ASKA Pharmaceutical Co.) for superovulation. At 14 hr after hCG injection, oocytes with cumulus cells (cumulus-oocyte complexes: COCs) were collected in 100 μL drops of HTF medium covered with paraffin oil. Cauda epididymal spermatozoa were collected from males and incubated in HTF medium for 2 hr for capacitation. Capacitated spermatozoa were added to each drop containing COCs at a final concentration of 2 × 10^5^ spermatozoa/mL. To remove cumulus cells, COCs were treated with hyaluronidase (1 mg/mL) for 5 min. At 8 hr after incubation (post-insemination), the formation of pronuclei was observed under a phase-contrast microscope (CKX41N-31PHP, Olympus, Tokyo, Japan). After IVF (Day 0), the embryos were cultured in KSOM medium until Day 4 (from 1-cell to blastocyst stages) and observed under the phase-contrast microscope.

### Mating test

Individual 8-week-old females were caged in pairs with individual 10-to 14-week-old males for 8 weeks. The numbers of litters and pups were counted in each cage.

### Statistical analysis

Statistical analysis was performed using GraphPad Prism 9. Data are represented as mean ± SD of three or more biological replicates. When comparing two groups, one-tailed or two tailed t*-* test was used, as indicated in the figure legends. To compare three or more groups, one-way ANOVA with the Tukey-Kramer *post hoc* test was used. The sample sizes represent independent biological replicates.

## Results

### Ovarian histology and AMH expression in AMH-TRECK Tg NOG mice

We previously established AMH-TRECK Tg mice on a C57BL/6 genetic background (Shinomura et al., 2014), which carry an *AMH* promoter-driven DTR-EGFP transgene (Figure 1A). We backcrossed these mice to NOG mice for at least eight generations to generate AMH-TRECK Tg NOG mice (Supplementary Figure S1A, B), which were fertile in both males and females. We detected no histological abnormalities in the ovaries of AMH-TRECK Tg NOG females (Figure 1B, C), consistent with previous observations in AMH-TRECK Tg C57BL/6 females (Shinomura et al., 2014). We then immunostained for AMH and showed that granulosa cells in primary and secondary follicles were strongly positive for AMH, whereas those in antral follicles were weakly positive or negative, and those in primordial follicles were negative for AMH in AMH-TRECK Tg NOG ovaries (Figure 1B, C). We also confirmed that GFP expression patterns in granulosa cells of AMH-TRECK Tg NOG ovaries were consistent with those of AMH expression patterns, although weak nonspecific fluorescent signals were detected in oocytes of both non-Tg and Tg ovaries (Figure 1D). All these ovarian characteristics in AMH-TRECK Tg NOG females were consistent with those previously reported in AMH-TRECK Tg C57BL/6 females (Shinomura et al., 2014). Based on these findings, all subsequent experiments reported here were conducted using AMH-TRECK Tg and non-Tg females on a NOG genetic background.

### Early effects of a single DT injection on postnatal AMH-TRECK Tg ovaries

We previously reported that a single DT injection at P3, when primary follicles containing AMH-positive granulosa cells first appear in the ovary, specifically depleted granulosa cells in developing follicles of AMH-TRECK Tg ovaries 4 days later (Shinomura et al., 2014). To further characterize the early effects of a single DT injection on the postnatal ovaries of AMH-TRECK Tg females, we performed a more detailed immunohistochemical and histological analysis. First, we injected a single dose of DT at P7, when primordial, primary, and secondary follicles are present (Kerr et al., 2006; Zhou et al., 2018), collected the ovaries 1 day later, and performed immunostaining for cleaved caspase-3, a marker of apoptosis (Figure 2A, B). At 1 day after a single DT injection, cleaved caspase-3-positive granulosa cells were rarely observed in non-Tg ovaries, whereas numerous positive cells were detected in both primary and secondary follicles of Tg ovaries. These cleaved caspase-3-positive follicles also contained granulosa cells exhibiting degenerating nuclear morphology. By contrast, granulosa cells in primordial follicles, oocytes at any follicular stage, and other somatic cell types were negative for cleaved caspase-3 in both non-Tg and Tg ovaries.

**Figure 2.**
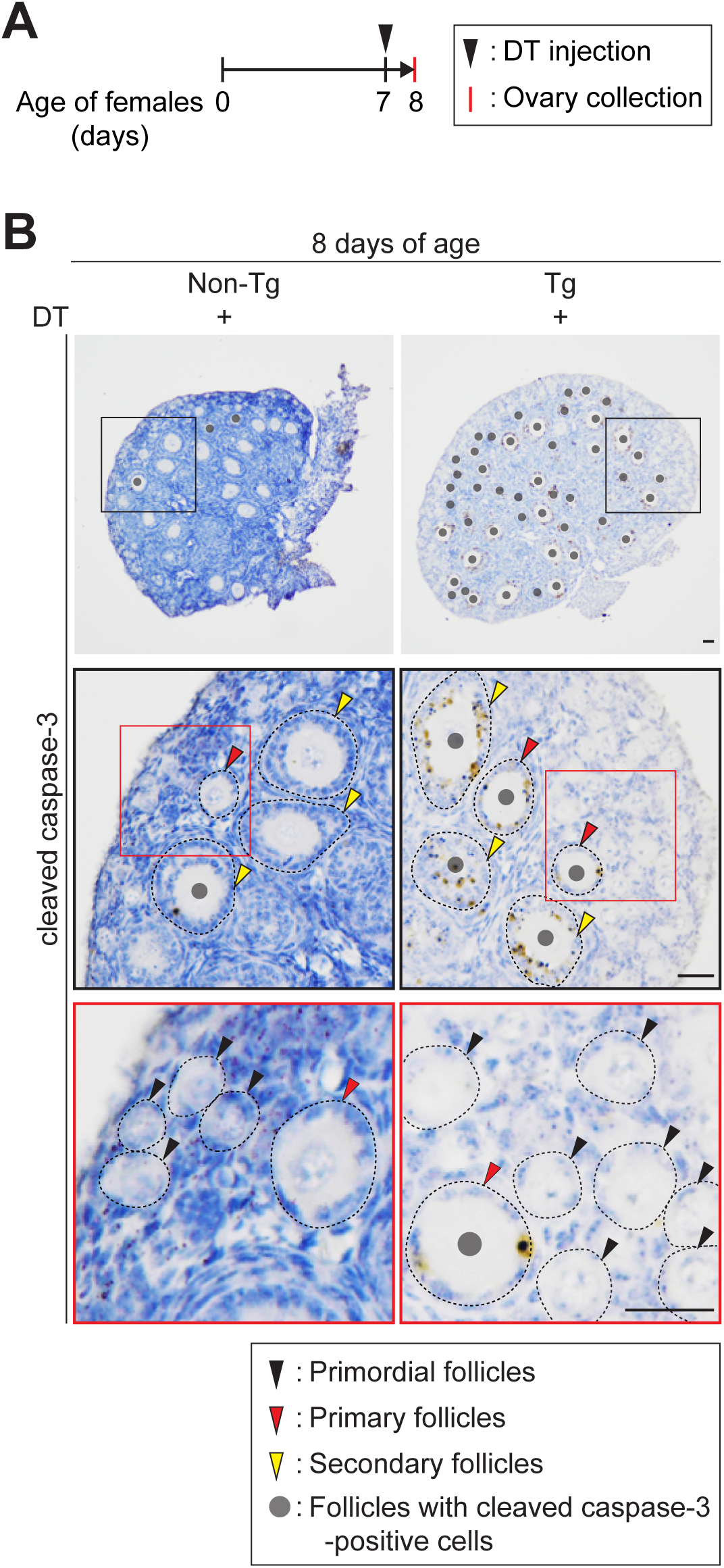
Single DT injection induces cleaved caspase-3-positive granulosa cells in primary and secondary follicles of postnatal ovaries in AMH-TRECK Tg NOG mice. (A) Experimental timeline of ovary collection following a single DT injection in postnatal female mice. Arrowhead, DT injection at postnatal day 7 (P7). Red bar, ovary collection at P8. (B) Ovary sections from non-Tg (left) and Tg (right) females at P8, immunostained for cleaved caspase-3 following a single DT injection at P7. Black boxed regions in the upper panels and red boxed regions in the middle panels are enlarged in the middle and lower panels, respectively. Arrowheads, primordial (black), primary (red), and secondary (yellow) follicles. Gray dots, follicles with cleaved caspase-3-positive cells. Scale bars, 25 μm.

Next, we injected a single dose of DT at P7, collected the ovaries 4 days later, and performed AMH immunostaining (Figure 3A, B). At 4 days after a single DT injection, most of the AMH-positive granulosa cells were depleted in AMH-TRECK Tg ovaries compared with non-Tg ovaries. In contrast, granulosa cells in primordial follicles were negative for AMH in both non-Tg and Tg ovaries. We then compared ovarian histologies between non-Tg and AMH-TRECK Tg ovaries (Figure 3C). At 4 days after a single DT injection, no discernible abnormalities were observed in granulosa cells or primary and secondary follicle structures in non-Tg ovaries. In AMH-TRECK Tg ovaries, most of the presumptive primary and secondary follicles, identified based on their sizes and positions within the ovary, exhibited a marked loss of granulosa cells surrounding the oocytes, whereas granulosa cells in primordial follicles remained unaffected.

**Figure 3.**
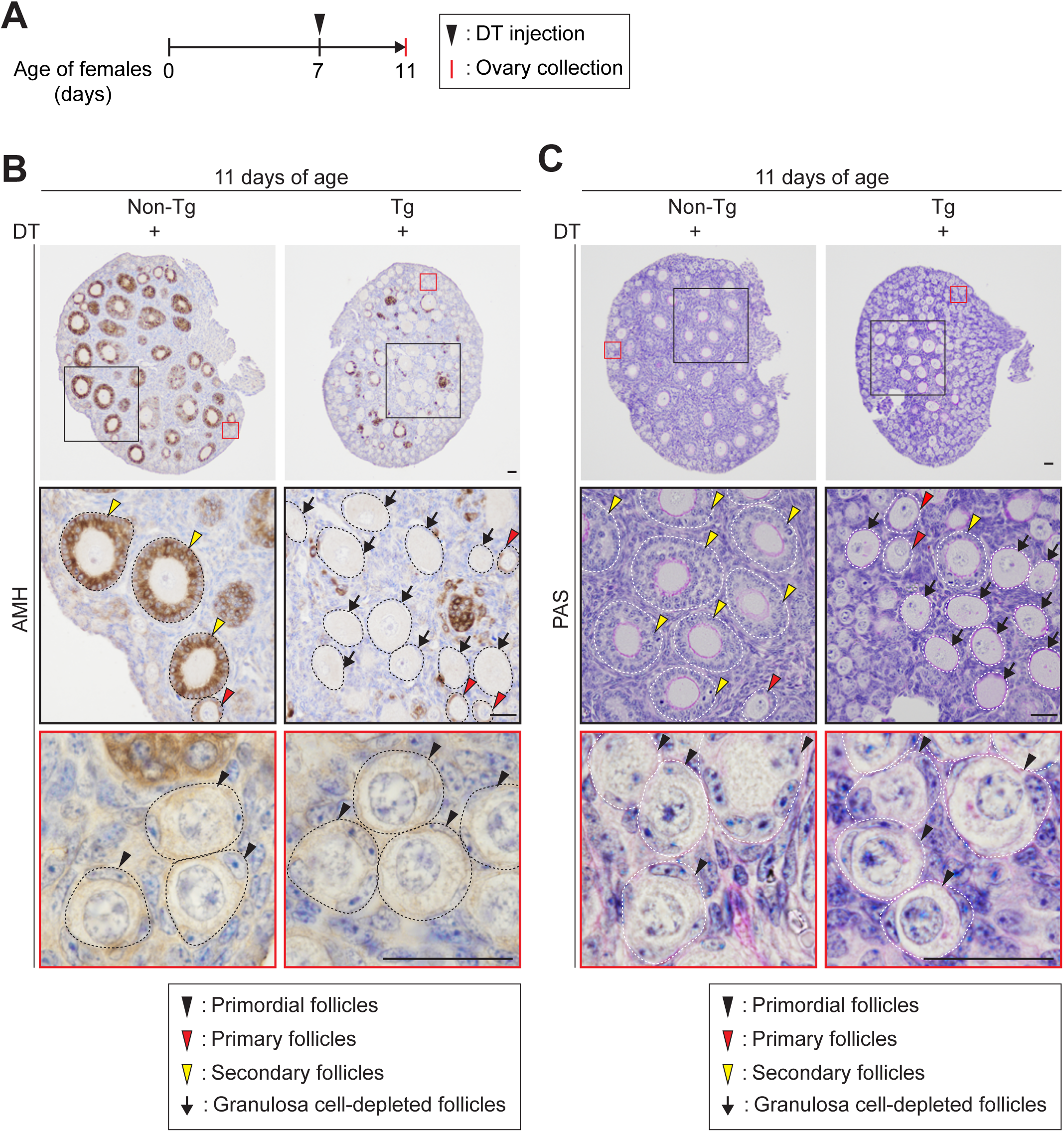
Single DT injection induces depletion of AMH-positive granulosa cells in primary and secondary follicles of postnatal ovaries in AMH-TRECK Tg NOG mice. (A) Experimental timeline of ovary collection following a single DT injection in postnatal females. Arrowhead, DT injection at postnatal day 7 (P7). Red bar, ovary collection at P11. (B, C) Ovary sections from non-Tg (left) and Tg (right) females at P11, immunostained for cleaved caspase-3 (B) or stained with hematoxylin and periodic acid-Schiff (PAS) (C) following a single DT injection at P7. Black and red boxed regions in the upper panels are enlarged in the middle and lower panels, respectively. Arrowheads, primordial (black), primary (red), and secondary (yellow) follicles. Arrows, granulosa cell-depleted follicles. Scale bars, 25 μm.

We also confirmed that oocytes within AMH-TRECK Tg ovaries showed no signs of depletion or degeneration 4 days after a single DT injection. We conclude that a single DT injection selectively depletes granulosa cells in primary and secondary follicles, but not those in primordial follicles, of AMH-TRECK Tg ovaries during postnatal ovarian development.

### Morphological and histological changes in adult AMH-TRECK Tg ovaries following repeated DT injections

We previously reported that a single DT injection at P3 depleted granulosa cells in developing follicles of AMH-TRECK Tg ovaries by P7 (Shinomura et al., 2014). By P28, however, AMH-positive primary and secondary follicles had been newly recruited from the AMH-negative primordial follicle pool (Shinomura et al., 2014). Because AMH inhibits the initiation of primordial follicle growth and adult *Amh* knockout (KO) mice exhibit a reduced primordial follicle pool (Durlinger et al., 2002; Durlinger et al., 2001; Durlinger et al., 1999), we hypothesized that repeated depletion of AMH-expressing granulosa cells should affect ovarian structure, including the number of primordial follicles, in adulthood. To test this prediction, we injected DT into mice weekly from 1 to 7 weeks of age and collected ovaries at 8 weeks of age (Figure 4A). Ovary sizes, weights, and weights per body weight were comparable between uninjected Tg and DT-injected non-Tg controls, whereas DT-injected Tg ovaries were smaller and had significantly reduced weights and weights per body weight compared with these controls (Figure 4B–D). Moreover, gross morphological examination revealed that large follicular structures observed in uninjected Tg and DT-injected non-Tg ovaries were absent from DT-injected Tg ovaries (Figure 4B).

**Figure 4.**
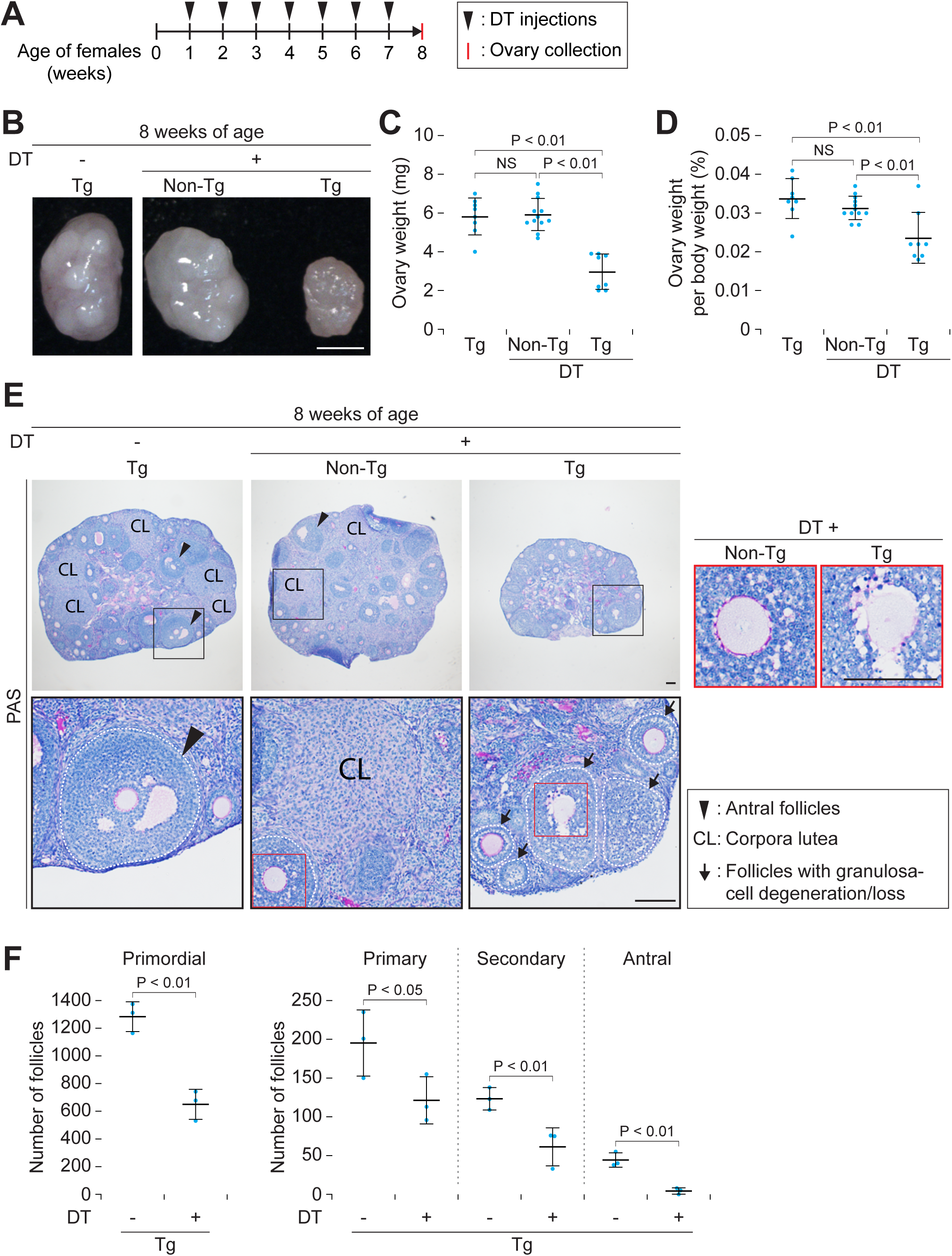
Effects of long-term repeated DT injections on adult ovaries in AMH-TRECK Tg NOG mice. (A) Experimental timeline of ovary collection following successive DT injections in females. Females received 7 DT injections at weekly intervals from 1 to 7 weeks of age. Arrowheads, weekly DT injections. Red bar, ovary collection at 8 weeks of age. (B) Gross morphology of ovaries at 8 weeks of age in Tg females without DT (left), non-Tg females with DT (middle), and Tg females with DT (right) injections. Scale bars, 1 mm. (C, D) Ovary weight (mg) (C) and ovary weight per body weight (%) (D) in Tg females without DT (n = 4), non-Tg females with DT (n = 6), and Tg females with DT (n = 4) injections at 8 weeks of age. Error bars, mean ± SD. Blue dots, individual ovaries. NS, not significant (*P* > 0.05); *P* < 0.01 (Tukey–Kramer test). (E) Ovary sections from Tg females without DT (left), non-Tg females with DT (middle), and Tg females with DT (right) injections at 8 weeks of age, stained with hematoxylin and periodic acid-Schiff (PAS). Black boxed regions in the upper panels are enlarged in the lower panels, and red boxed regions in the lower panels are enlarged in the far-right panels. Arrowheads, antral follicles. CL, corpora lutea. Arrows, follicles with granulosa cell degeneration and/or partial loss of granulosa cells. Dashed lines, basement membranes of follicles. Scale bars, 100 μm. (F) Number of primordial, primary, secondary, and antral follicles in Tg females without DT (n = 3) and with DT (n = 3) injections at 8 weeks of age. Error bars, mean ± SD. Blue dots, individual ovaries. *P* < 0.05, *P* < 0.01 (one-tailed t-test).

To further characterize ovarian structural changes, we examined ovarian histology among the three groups (Figure 4E). In uninjected Tg and DT-injected non-Tg controls, antral follicles and corpora lutea were observed in the ovaries at 8 weeks of age. In contrast, in DT-injected Tg ovaries, antral follicles and corpora lutea were rarely observed, and developing follicles contained degenerating granulosa cells and/or showed partial loss of granulosa cells (Figure 4E). We then counted the numbers of primordial, primary, secondary, and antral follicles in whole ovaries. Importantly, the numbers of all follicle types, including primordial follicles, were significantly reduced in DT-injected Tg ovaries compared with uninjected Tg ovaries (Figure 4F). We conclude that repeated depletion of AMH-expressing granulosa cells results in ovarian structural defects, including reduced numbers of primordial, primary, secondary, and antral follicles.

### Effects of repeated DT injections on ovulation, IVF, and early embryonic development in adult AMH-TRECK Tg females

We then tested whether repeated DT injections affect oocyte quality in AMH-TRECK Tg females. In DT-injected Tg ovaries, although antral follicles were rarely observed and developing follicles contained degenerating granulosa cells and/or showed partial loss of granulosa cells, late secondary follicles, identified based on their size and remaining granulosa cell layers, were present (Figure 4E, F). Because late secondary (preantral) follicles retain the capacity to respond to gonadotropin stimulation and produce viable ovulated oocytes (McGee and Hsueh 2000; Liu et al., 2002), we injected PMSG and hCG into adult AMH-TRECK Tg females following repeated DT injections (Figure 5A). At 14 hr after hCG injection, DT-injected Tg females showed a significantly reduced number of ovulated oocytes, and some of these oocytes exhibited partial denudation of surrounding granulosa/cumulus cells compared with DT-uninjected Tg controls (Figure 5B, C). Moreover, DT-injected Tg females showed a significantly reduced proportion of mature oocytes, identified based on first polar body extrusion, and a significantly increased proportion of degenerated or fragmented oocytes. In contrast, the proportion of immature oocytes was unchanged compared with DT-uninjected Tg controls (Figure 5D), indicating that the reduced proportion of mature oocytes was primarily due to the oocyte degeneration or fragmentation.

**Figure 5.**
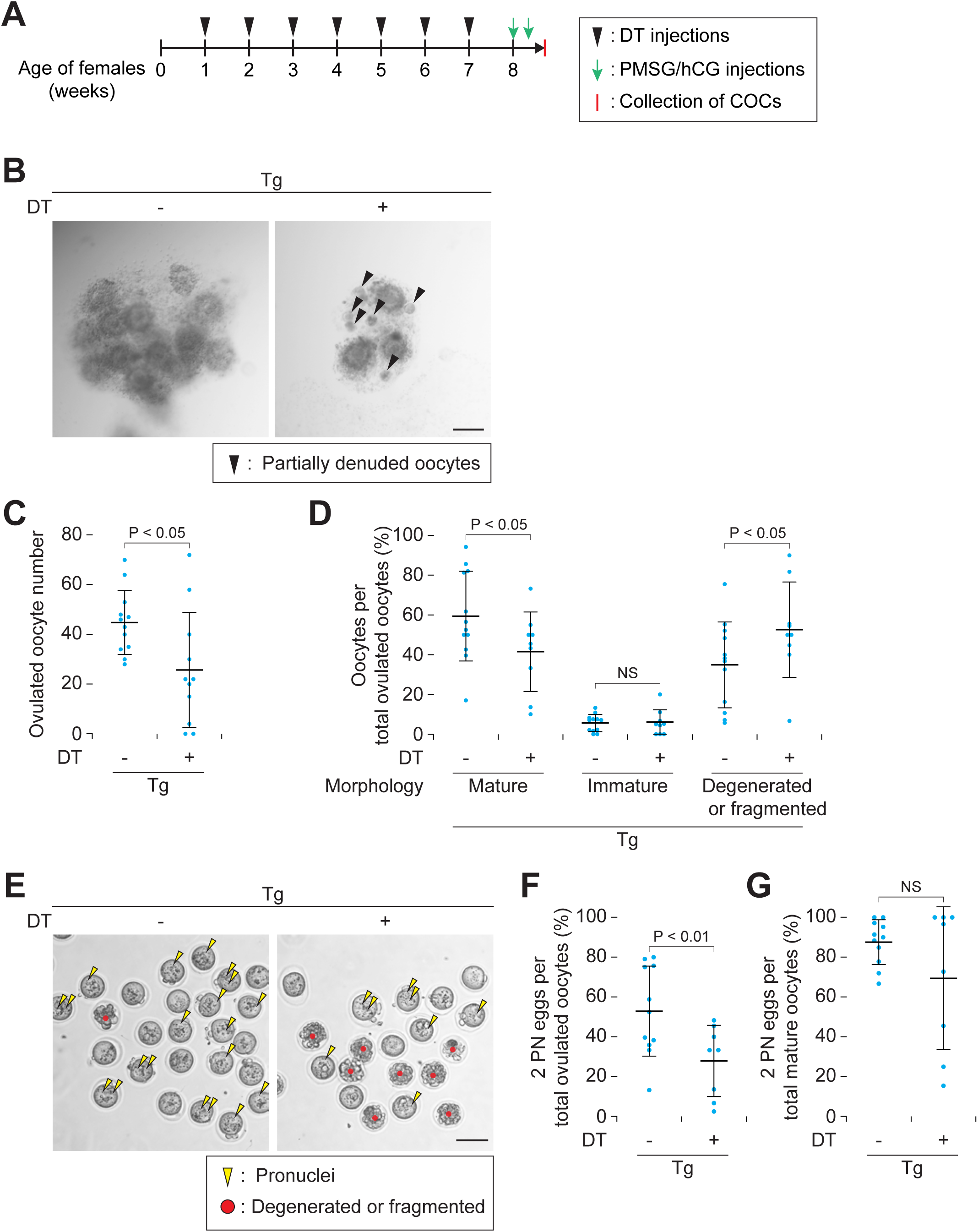
Effects of long-term repeated DT injections on oocytes and IVF in adult AMH-TRECK Tg NOG mice. (A) Experimental timeline of ovary collection following successive DT and hormone injections in females. Females received 7 DT injections at weekly intervals from 1 to 7 weeks of age, followed by PMSG and hCG for superovulation at 8 weeks of age. Arrowheads, weekly DT injections. Green arrows, pregnant mare serum gonadotropin (PMSG) and human chorionic gonadotropin (hCG) injections. Red bar, collection of oocytes with cumulus cells (cumulus-oocyte complexes: COCs) at 14 hr after hCG injection. (B) COCs collected from Tg females without (left) and with DT (right) injections. Arrowheads, partially denuded oocytes. Scale bars, 200 μm. (C) Ovulated oocyte number in Tg females without DT (n = 12) and with DT (n = 11) injections. Error bars, mean ± SD. Blue dots, total oocyte numbers from both ovaries of individual females. *P* < 0.05 (one-tailed t-test). (D) Mature, immature, and degenerated/fragmented oocytes per total ovulated oocytes (%) in Tg females without DT (n = 12) and with DT (n = 9) injections. Error bars, mean ± SD. Blue dots, individual females. NS, not significant (*P* > 0.05); *P* < 0.05 (one-tailed t-test). (E) Representative images of eggs at 8 hr after IVF from Tg females without (left) and with DT (right) injections. Ovulated COCs were co-incubated with capacitated spermatozoa for 8hr, after which cumulus cells were removed for imaging. Yellow arrowheads, visible pronuclei. Red dots, degenerated or fragmented eggs. Scale bars, 100 μm. (F, G) Two-pronuclear (2PN) eggs per total ovulated oocytes (%) (F) and per total mature oocytes (%) (G) in Tg females without DT (n = 11) and with DT (n = 8) injections. Error bars, mean ± SD. Blue dots, individual females. NS, not significant (*P* > 0.05); *P* < 0.01 (one-tailed t-test).

We then performed IVF to evaluate the fertilizability of oocytes ovulated from the DT-injected Tg ovaries. We co-incubated ovulated COCs with capacitated spermatozoa for 8hr (Figure 5E) and found that the number of fertilized eggs, identified by 2PN formation, per total ovulated oocytes was significantly lower in DT-injected Tg females than in uninjected Tg controls (Figure 5F). Importantly, the number of fertilized eggs per total mature oocytes were comparable between DT-injected Tg and uninjected Tg females (Figure 5G). These results indicate that the reduced fertilization rate of ovulated oocytes was mainly due to the increased number of degenerated or fragmented oocytes, and that mature oocytes in DT-injected Tg females retained the capacity to be fertilized.

To determine whether the fertilized eggs (embryos) derived from mature oocytes in DT-injected Tg females could develop normally, we cultured these embryos and monitored their development until the blastocyst stage (Day 4 after IVF). We found that 2-cell stage embryo rates at Day 1 were comparable between DT-injected Tg and uninjected Tg females (Figure 6A–B). At Day 4, blastocyst formation rates and hatching blastocyst rates were also comparable between DT-injected Tg and uninjected Tg females (Figure 6C–E), indicating that the fertilized eggs derived from mature oocytes in DT-injected Tg females can develop into blastocysts.

**Figure 6.**
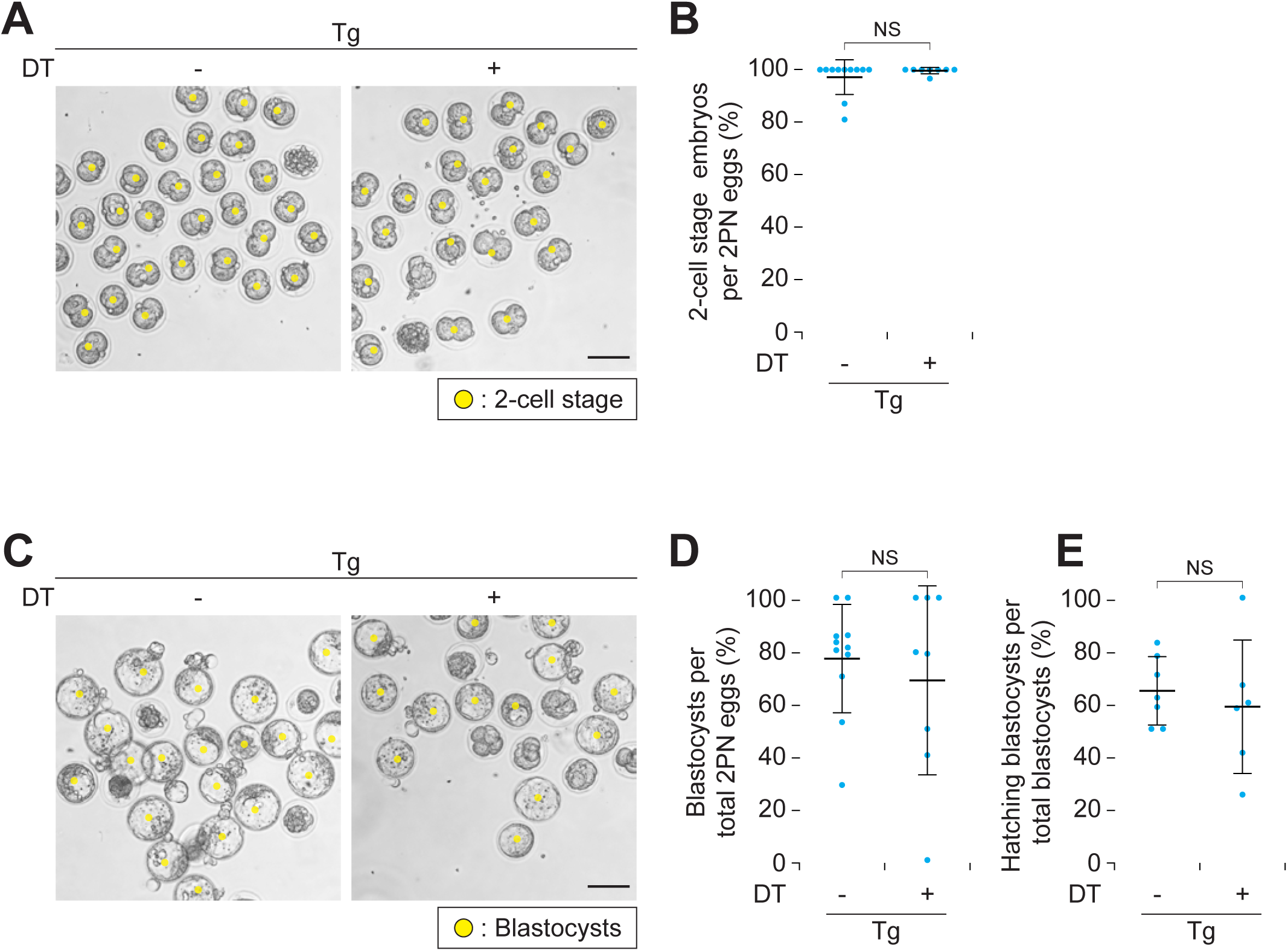
Effects of long-term repeated DT injections on early embryonic development following IVF in adult AMH-TRECK Tg NOG mice. (A) Eggs at 24 hr after IVF from Tg females without (left) and with DT (right) injections. Yellow dots, 2-cell stage embryos. Scale bars, 100 μm. (B) 2-cell stage embryos per total two-pronuclear (2PN) eggs (%) in Tg females without DT (n = 11) and with DT (n = 8) injections. Error bars, mean ± SD. Blue dots, individual females. NS, not significant (*P* > 0.05; one-tailed t-test). (C) Eggs at 96 hr after IVF from Tg females without (left) and with DT (right) injections. Yellow dots, blastocysts. Scale bars, 100 μm. (D, E) Blastocysts per total 2PN eggs (%) in Tg females without DT (n = 11) and with DT (n = 8) injections (D) and hatching blastocysts per total blastocysts (%) in Tg females without DT (n = 7) and with DT (n = 6) injections (E). Error bars, mean ± SD. Blue dots, individual females. NS, not significant (*P* > 0.05; one-tailed t-test).

### Effects of repeated DT injections on fertility of adult AMH-TRECK Tg females

We monitored *in vivo* fertility of repeated DT-injected adult AMH-TRECK Tg females by mating with non-Tg males for 8 weeks from 8 weeks of age (Figure 7A). In DT-injected Tg females, the number of litters, total pup numbers, and pup numbers per litter were significantly decreased compared with uninjected Tg females (Figure 7B–D). We conclude that repeated DT injections into AMH-TRECK Tg females from 1 to 7 weeks of age reduce their fertility.

**Figure 7.**
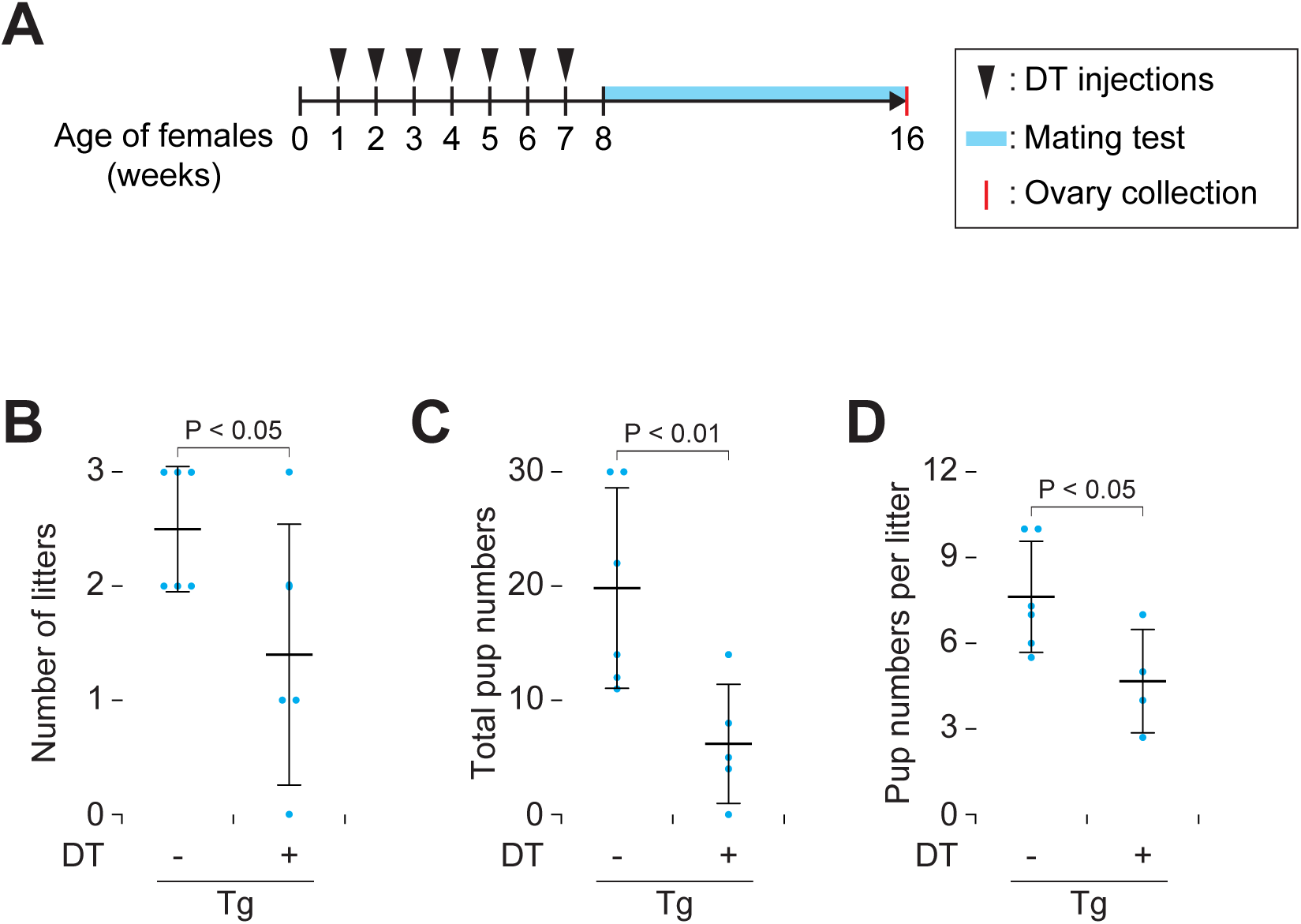
Long-term repeated DT injections reduce in vivo fertility in adult AMH-TRECK Tg NOG mice. (A) Experimental timeline of ovary collection following successive DT injections and a mating test in females. Females received 7 DT injections at weekly intervals from 1 to 7 weeks of age and were caged in pairs with a non-Tg male mouse from 8 to 16 weeks of age (blue box). Arrowheads, weekly DT injections. Red bar, ovary collection at 16 weeks of age. (B-D) Number of litters (B), total pup numbers (C), and pup numbers per litters (D) in Tg females without DT (n = 6) and with DT (n = 5) injections. One female in the DT-injected group produced no litter (B) and was therefore excluded from the number in (D). Error bars, mean ± SD. Blue dots, individual females. *P* < 0.05, *P* < 0.01 (one-tailed t-test).

### Recovery of folliculogenesis, but not primordial follicle numbers, after an 8-week interval following repeated DT injections in AMH-TRECK Tg females

We tested whether ovarian structure, including folliculogenesis and follicle numbers, recovers after an 8-week interval following repeated DT injections in AMH-TRECK Tg females. To address this question, we collected ovaries from DT-injected AMH-TRECK Tg females at 16 weeks of age after the mating test (Figure 7A). Both ovary weights and ovary weights per body weight were comparable between DT-injected Tg and uninjected Tg controls (Figure 8A, B). To characterize ovarian structures, we performed histological analysis of ovaries and found that DT-injected Tg ovaries at 16 weeks of age contained antral follicles and corpora lutea (Figure 8C), both of which were rarely observed at 8 weeks of age (Figure 4E). We then counted the numbers of primordial, primary, secondary, and antral follicles in whole ovaries at 16 weeks of age (Figure 8D). The number of primordial follicles was significantly lower in DT-injected Tg than in uninjected Tg controls, whereas the numbers of primary follicles, secondary follicles, and antral follicles were comparable between the two groups. We conclude that folliculogenesis and the numbers of primary, secondary, and antral follicles, but not those of primordial follicles, recover after an 8-week interval following repeated DT injection in AMH-TRECK Tg females.

**Figure 8.**
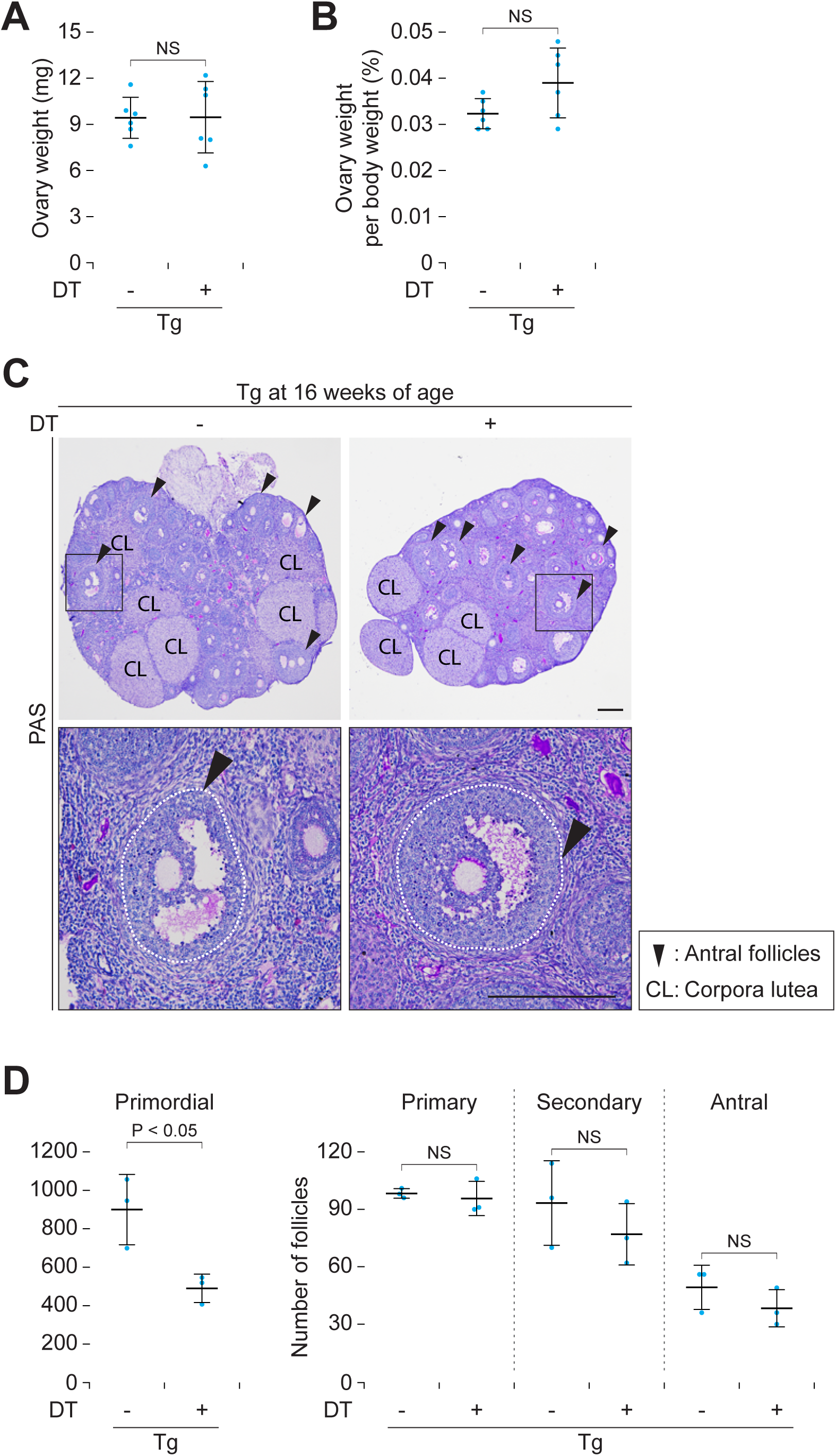
Follicular development, but not primordial follicle numbers, recovers after cessation of long-term repeated DT injections in adult AMH-TRECK Tg NOG mice. (A, B) Ovary weight (mg) (A) and ovary weight per body weight (%) (B) in Tg females without DT (n = 3) and with DT (n = 3) injections at 16 weeks of age following the experimental timeline shown in Figure 7A. Error bars, mean ± SD. Blue dots, individual ovaries. NS, not significant (*P* > 0.05; two-tailed t-test). (C) Ovary sections from Tg females without (left) and with DT (right) injections at 16 weeks of age, stained with hematoxylin and periodic acid-Schiff (PAS). Black boxed regions in the upper panels are enlarged in the lower panels. Arrowheads, antral follicles. CL, corpora lutea. Dashed lines, basement membranes of follicles. Scale bars, 200 μm. (D) Number of primordial, primary, secondary, and antral follicles in Tg females without DT (n = 3) and with DT (n = 3) injections at 8 weeks of age. Error bars, mean ± SD. Blue dots, individual ovaries. NS, not significant (*P* > 0.05); *P* < 0.05 (one-tailed t-test).

## Discussion

We investigated the comprehensive effects of AMH-expressing granulosa cell depletion on postnatal and adult ovaries, as well as female fertility, using AMH-TRECK Tg females. We found in Tg females that a single DT injection induces rapid apoptotic depletion of granulosa cells in developing follicles and that repeated DT injections result in ovarian structural changes and reduced fertility. Here we demonstrate that selective depletion of AMH-expressing granulosa cells *in vivo* impairs follicular development and fertility in female mice.

We found that, at 1 day after a single DT injection, both primary and secondary follicles of AMH-TRECK Tg ovaries contain granulosa cells positive for cleaved caspase-3 and/or exhibiting degenerating nuclear morphology. Because the characteristic nuclear morphological changes of apoptosis are known to follow activation of caspase-3 (Porter and Janicke 1999; He et al., 2009), our findings suggest that granulosa cell function are likely impaired within 1day of a single DT injection. By contrast, granulosa cells in primordial follicles, oocytes at any follicular stage, and other somatic cell types are negative for cleaved caspase-3 in Tg ovaries, and no depletion of these cells are observed at 4 days after a single DT injection. These findings are consistent with previous studies showing that mouse cells are naturally resistant to DT (Naglich et al., 1992; Mitamura et al., 1995; Cha et al., 2003). We thus conclude that a single DT injection selectively and conditionally depletes granulosa cells of primary and secondary follicles within 4 days in postnatal ovaries of AMH-TRECK Tg females.

After repeated DT injections weekly from 1 to 7 weeks of age in AMH-TRECK Tg females, antral follicles and corpora lutea are rarely observed at 8 weeks of age, indicating that granulosa cell loss impairs follicular development and subsequent ovulation. This impaired follicle development is further supported by our findings that the numbers of primary and secondary follicles are decreased and that these developing follicles contain degenerating granulosa cells and/or exhibit partial loss of granulosa cells. Given that a single DT injection at P7 results in severe loss of granulosa cells in developing follicles 4 days later, the partial loss of granulosa cells at 8 weeks of age may reflect the weekly intervals between DT injections and the timing of ovary collection (7 days after the final DT injection). At 4 days after DT injection to P7, a few AMH-expressing granulosa cells remain in some developing follicles of Tg ovaries. These residual AMH-expressing granulosa cells after DT injection may proliferate and partially repopulate developing follicles in adult ovaries, as granulosa cells undergo extensive proliferation during follicular development (McGee and Hsueh 2000; Edson et al., 2009).

In AMH-TRECK Tg ovaries at 8 weeks of age, repeated DT injections decrease not only developing follicle numbers but also primordial follicle numbers, suggesting that repeated depletion of AMH-expressing granulosa cells leads to a reduction in the primordial follicle pool. Our findings are consistent with previous studies demonstrating that AMH inhibits the transition of primordial follicles to developing follicles and that *Amh* KO mice exhibit a reduced primordial follicle pool (Durlinger et al., 1999; Durlinger et al., 2001; Durlinger et al., 2002).

However, in *Amh* KO mice, a reduction in the number of primordial follicles becomes evident at 4 months of age, and reductions in developing follicles are observed at 13 months of age (Durlinger et al., 1999; Durlinger et al., 2001; Durlinger et al., 2002). Thus, depletion of AMH-expressing granulosa cells may affect ovarian follicle development at an earlier time point and/or more broadly than loss of the *Amh* gene alone. Future studies are needed to determine whether depletion of AMH-expressing granulosa cells alters the ovarian microenvironment, including interfollicular interactions, as well as the signaling pathways that regulate primordial follicle activation and follicular development.

Following PMSG and hCG stimulation, repeated DT-injected adult AMH-TRECK Tg females exhibit a reduced number of ovulated oocytes, and some ovulated oocytes show partial loss of surrounding granulosa/cumulus cells. These findings align with ovarian histological data showing reduced numbers of developing follicles and partial loss of granulosa cells. In ovulated oocytes, the proportion of mature oocytes decreases, accompanied by an increase in the proportion of degenerated or fragmented oocytes. By contrast, the proportion of immature oocytes remains unchanged, suggesting that repeated depletion of AMH-expressing granulosa cells does not primarily cause oocyte maturation arrest but rather affects oocyte viability.

Moreover, mature oocytes from repeated DT-injected Tg ovaries retain the capacity for *in vitro* fertilization and blastocyst development, suggesting that reduced *in vivo* fertility is largely due to the reduced number of ovulated oocytes and the lower proportion of mature oocytes.

We found that, after an 8-week cessation of DT treatment in adult AMH-TRECK Tg females, ovarian weight and follicular development recover. Specifically, these Tg ovaries at 16 weeks of age exhibit recovery of antral follicles and corpora lutea, which are rarely observed in repeated DT-injected Tg ovaries at 8 weeks of age, whereas the primordial follicle pool remains reduced. Our findings provide evidence that the primordial follicle pool retains the capacity for follicular development even after repeated depletion of AMH-expressing granulosa cells and reduction of developing follicles in the ovaries. Importantly, although the number of primordial follicles remains reduced, the numbers of primary, secondary, and antral follicles in DT-injected Tg ovaries are restored to levels comparable to those in DT-uninjected Tg ovaries at 16 weeks of age. This finding suggests that the number of primary follicles recruited from primordial follicles is not directly determined by the total size of primordial follicle pool in the ovary. It will be of great interest to determine the mechanisms underlying the recruitment of a precise number of developing follicles in the ovary.

*In vitro* culture systems using isolated oocytes and granulosa cells, as well as follicle culture, have provided valuable insights into local cellular interactions (Schroeder and Eppig 1989; Emori and Sugiura 2014; Telfer et al., 2023). Complementary *in vivo* approaches are important for understanding complex follicular dynamics involving interfollicular communication and the ovarian microenvironment. To date, numerous *in vivo* studies have largely relied on gene disruption approaches using KO model, such as *Amh*, *Fshb*, *Foxo3*, and *Bcl2* κΟ mice (Ratts et al., 1995; Durlinger et al., 2001; Castrillon et al., 2003). However, the effects on granulosa cells and follicles observed in these KO mice are secondary consequences of gene disruption. Thus, direct and conditional depletion of granulosa cells using AMH-TRECK Tg females enables the assessment of their primary effects *in vivo*. This model may also provide insights into mechanisms underlying ovarian reserve depletion observed in human primary ovarian insufficiency (POI) (; Nelson 2009; Federici et al., 2024). Here, we generated and analyzed AMH-TRECK Tg female mice on a NOG background, which may provide a useful platform for future transplantation-based studies in females. Specifically, allogeneic transplantation between different mouse strains and/or xenogeneic transplantation into DT-injected Tg females may help determine how transplanted mouse or human granulosa cells, or ovarian tissues, influence follicular development through secreted factors.

## Author contributions

T.E., R.T., A.T., Y.K., and M.K.A. designed the research; T.E., M.T., K. H., F.O., N.O., K.W., T.L., Y.N., Y.F., M.G., and R.T. performed the research; T.E., M.T., K.H., F.O., N.O., K.W., M.G., and R.T. analyzed the data; K.S., Y.H., and N.M. provided advice on study design and data interpretation; and T.E., Y.K., and M.K.A. wrote the paper.

## Acknowledgements

We thank the Center for Experimental Animals, Institute of Science Tokyo, especially Hitomi Takahashi, for animal care and technical support, Dr. Hiroshi Suemizu, Dr. Hiroshi Yomogita, Dr. Hinako Takase, and Dr. Chihiro Emori for helpful discussion or technical assistance, and Maaya Sada for critical reading of the manuscript.

