## Supplemental Figure S1 for "Selective depletion of AMH-expressing granulosa cells *in vivo* impairs follicular development and fertility in female mice"

### Supplementary Figure S1

**A**

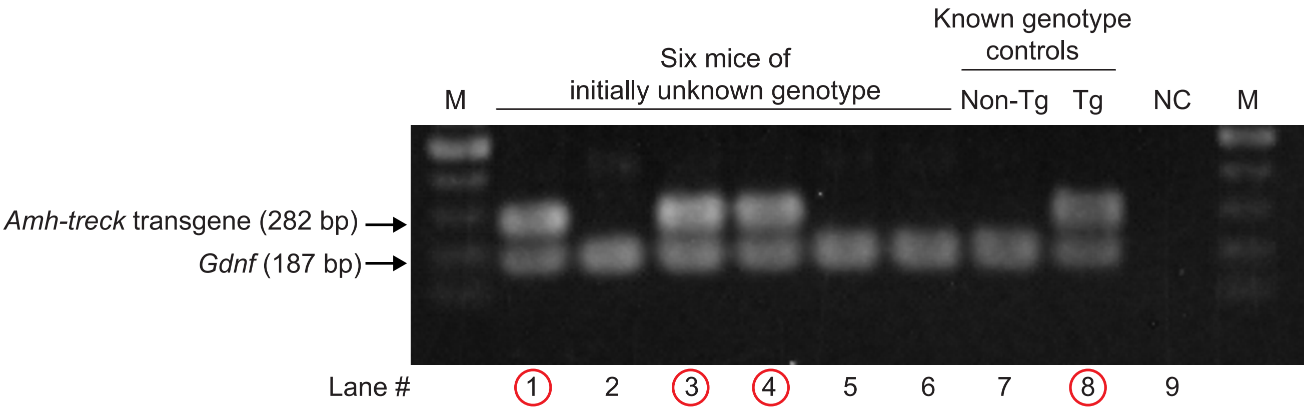


**B**

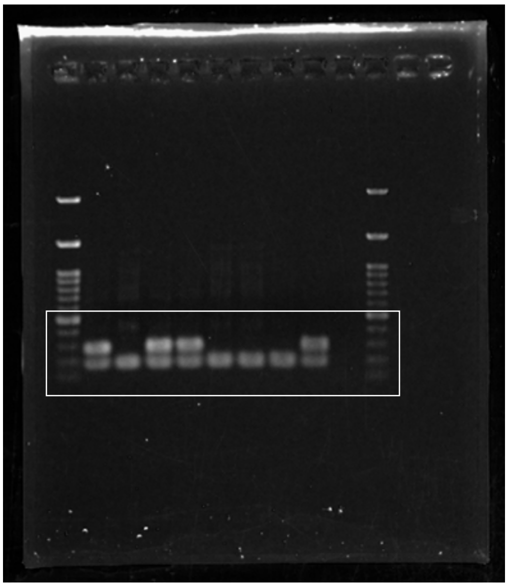

**Supplementary Figure S1. Representative genotyping PCR for identification of AMH-TRECK Tg and non-Tg mice.**

(A) Multiplex PCR was performed using primers specific for the *AMH*-TRECK transgene (282 bp) and *Gdnf* (187 bp) as an internal control. The positions of the primers for *AMH*-TRECK transgene are shown in Figure 1A. The gel includes six mice of initially unknown genotype (lanes 1–6) and known non-Tg and Tg mice used as genotype controls (lanes 7 and 8, respectively). NC, negative control (no template). M, 100-bp DNA ladder markers. Red circles, Tg mice. (B) Original uncropped gel image. The white boxed region corresponds to the cropped area shown in (A).
